# *CREB3* gain of function variants protect against ALS

**DOI:** 10.1101/2024.10.10.617542

**Authors:** Salim Megat, Christine Marques, Marina Hernan Godoy, Chantal Sellier, Geoffrey Stuart-Lopez, Sylvie Dirrig-Grosch, Charlotte Gorin, Aurore Brunet, Mathieu Fischer, Céline Keime, Pascal Kessler, Marco Antonio Mendoza-Parra, Sonja Scholz, Luigi Ferrucci, Albert Ludolph, Bryan Traynor, Adriano Chio, Luc Dupuis, Caroline Rouaux

**Author notes:** Co-corresponding authors &. These authors contributed equally to this work (Salim Megat, Christine Marques).

## Abstract

Amyotrophic lateral sclerosis (ALS) is a fatal and rapidly evolving neurodegenerative disease that arises from the loss of glutamatergic corticospinal neurons (CSN) and cholinergic motoneurons (MN). The disease is mostly sporadic, but genetics is expected to highly contribute to disease onset and progression. Genome wide association studies identified a few genetic disease modifiers, mostly associated with a negative outcome, and demonstrated that ALS is primarily a disease of excitatory glutamatergic neurons. Here, we reasoned that at least a subpart of genetic disease modifiers may directly modulate the molecular pathways selectively activated in vulnerable neurons as the disease progresses, and concentrated on CSN for their selective vulnerability and glutamatergic identity. We implemented comparative cross-species transcriptomics using snRNAseq data from postmortem motor cortex of ALS patients and controls, and longitudinal RNAseq data from anatomically defined CSN purified from the *Sod1^G86R^* mouse model of ALS. We report that disease vulnerable neuronal populations undergo ER stress and altered mRNA translation, and identify the transcription factor CREB3 and its regulatory network as a resilience marker of neuronal dysfunction in ALS. Using genetic and epidemiologic analyses we further identify the rare variant CREB3^R119G^ (rs11538707) as a new disease modifier in ALS. Through gain of function, CREB3^R119G^ decreases both the risk of developing ALS and the progression rate of ALS patients. This study reveals novel genetic variants that protect against ALS and highlights the benefice of combining transcriptomics and genetics to identify new disease modifiers and therapeutic targets.

## Introduction

Amyotrophic lateral sclerosis (ALS) is a fatal neurodegenerative disease that manifests as a progressive muscular weakness and paralysis, leading to death within only two to three years upon symptom onset ^1^. ALS results from the progressive degeneration of two neuronal populations involved in voluntary motor control: the glutamatergic corticospinal neurons (CSN, or upper motor neurons) in the motor cortex, and the cholinergic motoneurons (MN or lower motoneurons), in the brainstem and spinal cord ^1^.

Whereas the vast majority of patients are considered sporadic, about 10% have a familial history, and causative genes have been identified in more than half of these cases, the most represented being *C90RF72, SOD1, TARDBP* and *FUS* ^2^. While ALS genetics is complex, causative mutations remarkably converge on recurrent dysregulated pathways including DNA repair, RNA and protein metabolism, intracellular trafficking and mitochondrial dysfunction ^2^. This suggests that at least a subgroup of genetic disease modifiers may modulate, either positively or negatively, the same cellular pathways that accompany the degeneration of vulnerable neurons. The most recent and largest GWAS to date identified 15 ALS-associated risk loci with enriched expression in excitatory glutamatergic neurons ^3^, a cell-type specificity that we ^4^ and others ^5^ recently confirmed. We thus reasoned that unravelling the deregulated cellular pathways that accompany CSN degeneration during the course of ALS may inform on putative new genetic disease modifiers and therapeutic targets. Pioneering snRNAseq of post-mortem motor cortex from patients and controls recently provided molecular access to vulnerable cortical excitatory neurons and offered valuable snapshots of these neuronal populations at disease end-stage ^5,6^. Yet, the population of CNS is altogether under-represented in the healthy human brain and further depleted during the course the disease ^5^, and its access limited to post-mortem time point further restrict our ability to understand their disease trajectory and means to cope with disease progression.

To circumvent this limitation, we employed cross-species transcriptomics, combining snRNAseq data from ALS patients, frontotemporal dementia (FTD) patients, and healthy individuals ^5^ to longitudinal RNAseq data of CSN purified from the motor cortex of a mouse model of ALS that recapitulates their progressive degeneration ^7^. We hypothesized that a conserved signature of CSN degeneration exists across species, and reasoned that the comparative integration of mouse and human transcriptomics could improve access to disease-relevant neuronal populations.

We report that comparative transcriptomics identified disease-vulnerable human neuronal populations in ALS patients, that primarily undergo ER stress and altered mRNA translation. We further identified the transcription factor CREB3 and its regulatory network as a resilience marker of neuronal dysfunction in human and mice. Finally, through genetic analyses, we identified a missense rare variant of *CREB3* (rs11538707, p.Arg119Gly), which, through gain of function mechanisms, confers a 40% reduction in the risk of developing ALS and is associated with a slower disease progression rate in ALS patients, extending disease duration by 12 months.

## Results

### Cross-species RNAseq integration prioritizes disease-vulnerable human neuronal populations

To unequivocally identify the disease vulnerable CSN amongst human neuronal populations, we integrated the snRNAseq dataset obtained by Pineda and collaborators from 23 sporadic and familial ALS patients, 26 FTLD patients and 17 aged-matched controls ^5^, to the cross-species snRNAseq dataset generated by Bakken and collaborators that allowed consensus classification of cell types ^8^ (**Fig. 1a**). Our analysis yielded a total of 245,143 nuclei integrated across the independent datasets allowing the identification of a total of 116 clusters (**Fig. 1b**). These were further hierarchically organized based on transcriptomic similarities and associated with their positioning across the cortical layers ^8^, providing a full representation of glutamatergic excitatory neurons, GABAergic inhibitory neurons, and non-neuronal cells (**Supplementary Fig. 1**). Noteworthily, the population of extra-telencephalic neurons from the cortical layer 5, L5-ET, suspected to represent ALS-vulnerable human CSN, yielded the smallest number of neurons (∼1000 neurons, corresponding to 0.6% of the total nuclei and 1.7% of all excitatory neurons), and displayed the lowest number of expressed genes compared to other excitatory neuron populations (**Supplementary Table 1**). More importantly, we observed that the gene-level dispersion was much higher for low to medium expressed genes and negatively correlated with the number of cells sequenced per individual (**Supplementary Fig. 2**). Together, this indicates that rare cell types, such as degenerating CSN, may hinder differential expression analysis and discovery of new potential target genes.

**Figure 1:**
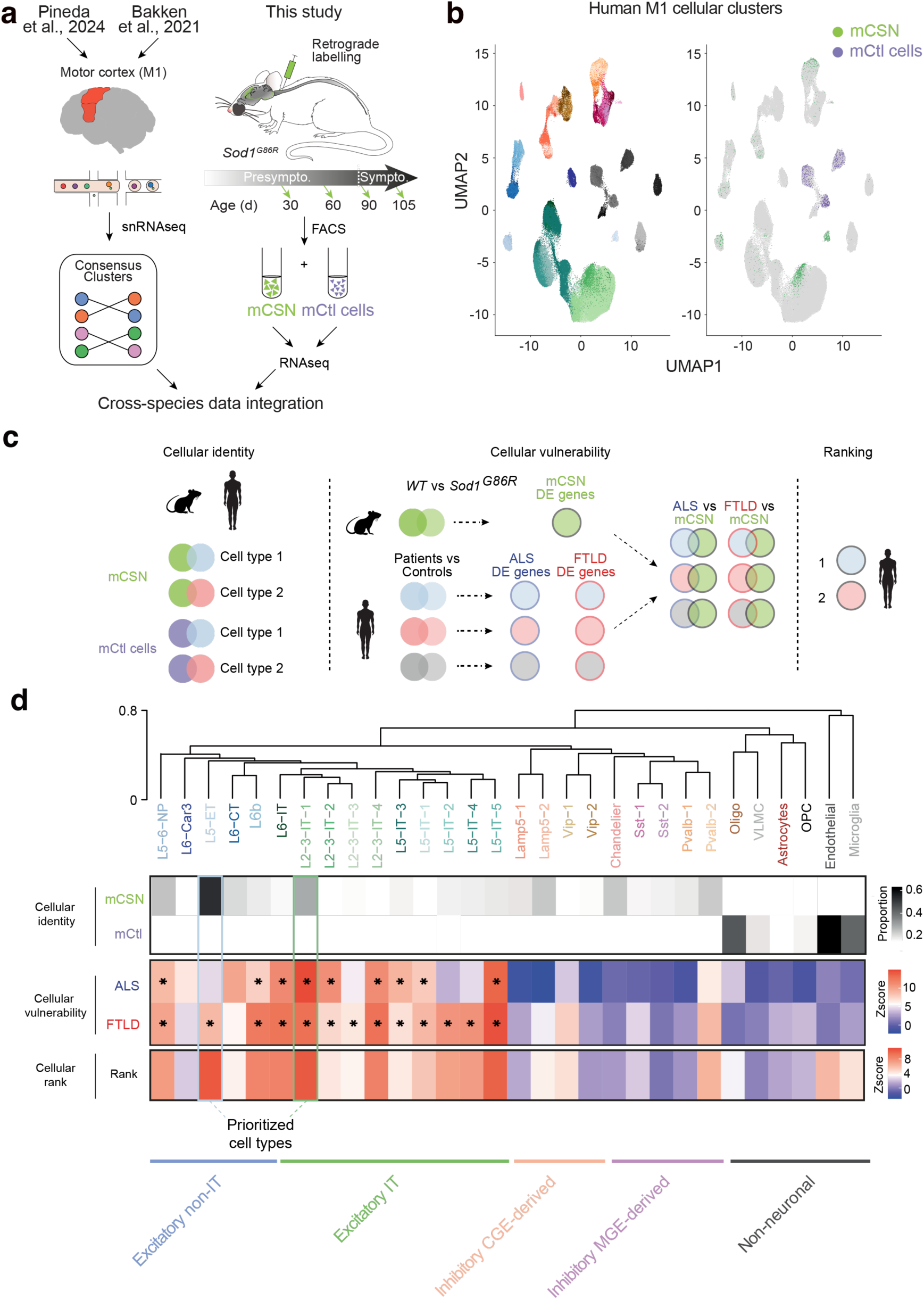
Cross-species transcriptomic analysis prioritizes vulnerable cell-types in ALS. **a.** Experimental design displaying integration of single nuclei RNAseq from ALS patients and healthy controls ^5^, the human motor cortex ^8^ and our database of mouse retrogradelly labelled CNS and control cells (mCSN and mCtl cells) from WT and *Sod1^G86R^* mice. **b.** T-SNE plot of the 116 clusters identified in the human M1 motor cortex (left) and expression of the mCNS and mCtl cells enriched genes in each cluster (right). **c.** Experimental design of the cross-species DEG enrichment analysis wherein DEG identified in mCSN were intersected with each of the 30 DEG cell populations in human ALS, FTD and healthy controls. **d.** Cluster dendrogram and heatmaps showing enrichment of mCSN and mCtl cells geneset in each of the 30 DE cell populations as well as enrichment of mCSN DEG in all DE cell populations. We observed a significant enrichment of the mCSN DEG in human excitatory neurons (Hypergeometric test: Bonferroni’s adjusted *p < 4^e-04^). Based on mCSN geneset enrichment and cross-species DE analysis, a cumulative score was calculated for each DE group and ranked to prioritize the most affected cell populations in ALS patients.

To identify presumable CSN among the different human neuronal populations, we generated a new dataset of anatomically-identified, disease-vulnerable mouse corticospinal neurons (mCSN) for comparison with the human dataset. mCSN were retrogradelly labelled from the cervical dorsal funiculus of *Sod1^G86R^* mouse model of ALS and their WT littermates, and purified at presymptomatic (30 and 60 days), and symptomatic (90 and 105 days) ages (**Fig. 1a**). Non-labelled cells from L2/3 were also purified from the same animals to serve as a control cellular population (mCtl cells) (**Fig. 1a**). Unsupervised hierarchical clustering separated mCSN from mCtl cells (**Supplementary Fig. 3a**) and indicated that a larger part of the variance observed between WT and *Sod1^G86R^* mice is attributable to mCSN (**Supplementary Fig. 3b**). We reasoned that cross-species RNAseq integration could be employed to *i)* identify the human neuronal populations that more closely resemble mCSN, independently of the disease condition or genotype (*i.e.* ‘cellular identity’, **Fig. 1b,c**), *ii)* identify the human neuronal populations whose differential gene expression (DGE) in ALS or FTLD conditions mimics mCSN DGE between *Sod1^G86R^*and control mice, (*i.e.* ‘cellular vulnerability’, **Fig. 1b,c**), and *iii)* prioritize human neuronal populations based on the combination of ‘cellular identity’ and ‘cellular vulnerability’ (*i.e.* ‘ranking’, **Fig. 1b,c**).

We first selected the top differentially expressed genes (DEG) between mCSN and mCtl cells (**Supplementary Table 2**) to create a mCSN geneset and its homologous human geneset (**Supplementary Table 3**) ^8^, integrated human and mouse datasets, and identified mCSN geneset in L5-ET (**Fig. 1b**), L5/6 NP, L5-IT and L2/3-IT neurons (**Supplementary Fig. 1**). mCtl cells genes instead were enriched non-neuronal populations: endothelial cells, microglia, and oligodendrocytes (**Supplementary Fig. 1**). To further maximize power with differential expression analysis, we used aggregated hierarchical clustering to regroup the initially identified 116 cell clusters into 30 cell populations. This confirmed that the mCSN geneset is expressed in a large proportion of L5-ET (∼60%) and a significant proportion (∼40%) of L2/3 IT-1 neurons (**Fig. 1d**). Importantly, these neurons express high amounts of *NEFH*, encoding the heavy subunit of neurofilament (**Supplementary Fig. 4**), typical of long-range projection neurons. This also confirmed the identity of mCtl cells as non-neuronal populations (**Fig. 1d**). Together, this shows that several sub-populations of excitatory neurons from the human motor cortex express the gene signature of mCSN, including L5/6 NP, L5-IT and L2/3-IT populations, in addition to the expected L5-ET neurons.

Second, we intersected DEG between WT and *Sod1^G86R^* mCSN (**Supplementary Table 4**) with DEG between controls and ALS patients, or DEG between controls and FTLD patients for each of the 30 cell populations (**Fig.1d, Supplementary Table 5 and 6**). This analysis confirmed a significant enrichment of mCSN DEG in human excitatory neuron populations as opposed to inhibitory and non-neuronal cell populations (**Supplementary Table 7 and 8**, Bonferroni’s adjusted *p < 4^e-04^). More precisely in ALS, this identified L5/6 NP, L6b, L6 IT, L 2/3 IT 1, 2 and 4, and L5 IT 1, 3 and 5 (**Fig. 1d**). The same populations were identified in FTLD, together with L5 ET, L2/3 IT 3, and L5 IT 2 and 4 (**Fig. 1d**).

Third, we ranked each human cell population based on the combination of the ‘cellular identity’ and ‘cellular vulnerability’ results, which prioritized L5-ET and L2/3 IT-1 neurons as the human populations that more closely recapitulate the transcriptomic signature of diseased mCSN (**Fig. 1d**). In whole, cross-species RNAseq integration allowed us to prioritize L5-ET and L2/3 IT-1 populations for further analyses.

### WGCNA consensus on prioritized cell populations reveals conserved gene regulatory network in ALS

We next aimed at unravelling disease-associated pathways and employed weighted-gene co-expression network analysis (WGCNA) consensus across species and prioritized cell types (**Fig. 2a**). WGCNA analysis identified four mRNA modules significantly correlated with the genotype (mice) or disease condition (humans) and labelled as lightyellow, darkgreen, turquoise and darkgrey modules according to the WGCNA conventions (Benjamini-Hochberg p < 0.05; **Fig. 2b, Supplementary Table 9**). We then constructed a graphical network of significantly associated modules to highlight hub genes, related weight of the genes within the network and connectivity (**Fig. 2c**), that illustrates the independent and relative proportions of non-overlapping networks. Assessing the conservation across species further allowed us to prioritize the turquoise and lightyellow modules that respectively showed strong (Zscore^preservation^ > 10) and moderate preservation (5 < Zscore^preservation^ < 10, **Fig. 2d**).

**Figure 2:**
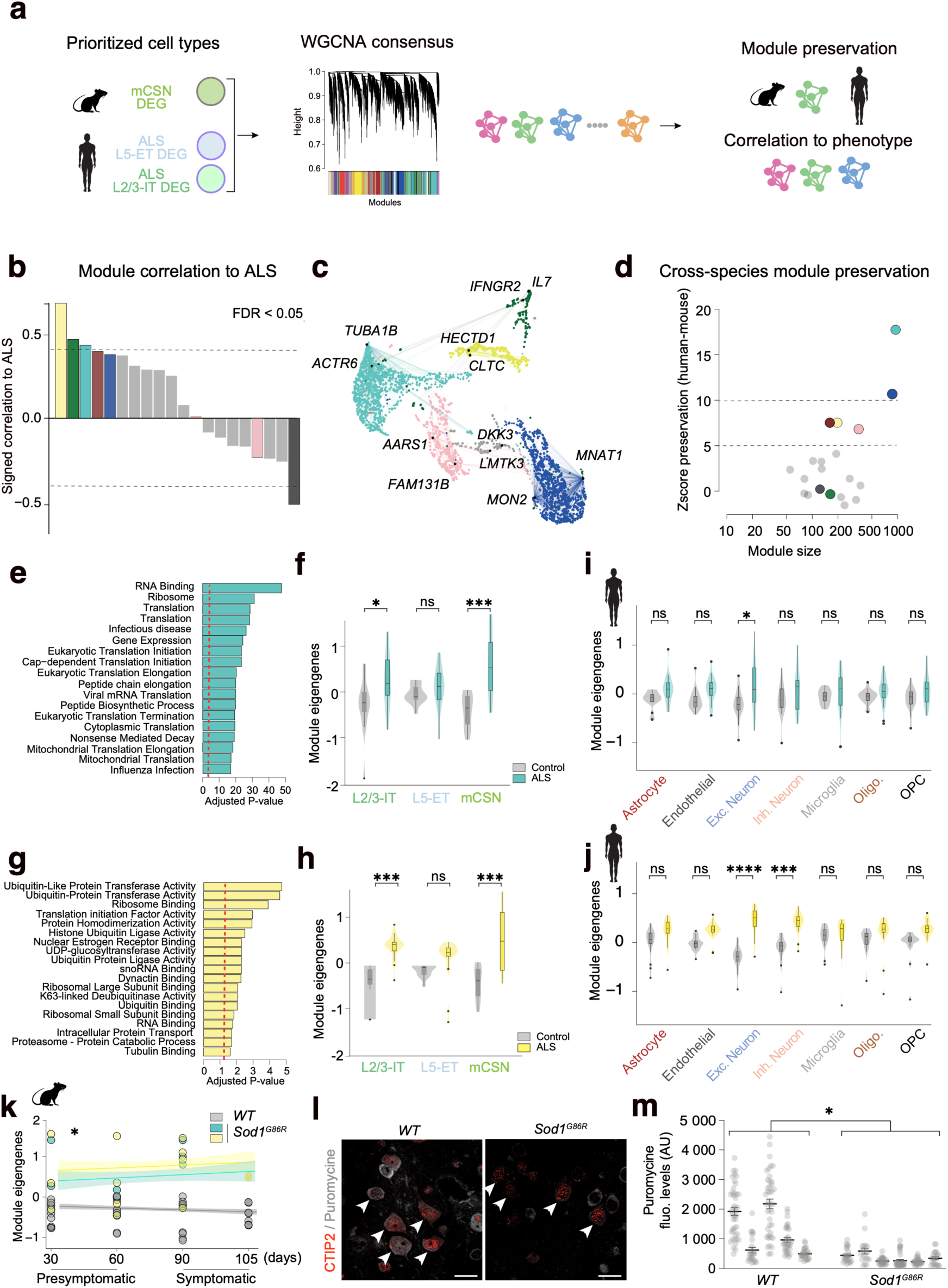
Cell-type specific consensus WGCNA identifies conserved regulatory networks. **a.** Experimental design showing the cross-species consensus WGCNA approach used on the prioritized cell types identified in Fig.1. **b.** Histogram showing conserved signed correlation of twenty WGCNA modules to ALS in mCSN and human L2/3-IT1 and L5-ET. Correlation p-value for each module were corrected using the Benjamini-Hochberg method. Dashed line represents WGCNA module with signed correlation FDR < 0.05. **c.** UMAP gene network of the lightyellow, turquoise, pink, blue and darkgrey modules and 2 hub genes per module. Network shows intra-module connectivity with each dot representing individual genes. The size of the dot is proportional to the importance of each gene (signed kME) in the network. **d.** Dot plot showing cross-species conservation of significant modules, with Zscore > 10 (turquoise and blue modules) considered to have strong preservation and 5 < Zscore < 10 to have moderate preservation (darkred, lightyellow and pink modules). **e.** GO-term enrichment analysis in the significant turquoise module with genes having signed kME (>0.6) in the network. Histograms represent significant terms above the red dashed line (FDR < 0.05). **f.** Violin plot showing module eigengenes expression of the turquoise module in human L2/3-IT1 and L5-ET and mCNS (ANOVA: F_interaction_ group*cell =4.88, **p<0.01) with a significantly increased expression in patients’ L2/3-IT1 neurons (Tukey-multiple comparisons: *p-adjusted=0.037) and *Sod1^G86R^* mCNS neurons (Tukey-multiple comparisons : ***p-adjusted<0.001) compared to healthy controls or WT animals respectively, and a trend to an increase in patients’ L5-ET (Tukey-multiple comparisons: p>0.05). **g.** GO-term enrichment analysis in the significant lightyellow module with genes having signed kME (>0.6) in the network. Histograms represent significant terms above the red dashed line (FDR < 0.05). **h.** Violin plot showing module eigengene expression of the lightyellow module in all three cell types (ANOVA: F_interaction_ group*cell=5.007, **p<0.01) with significantly increased expression in patients’ L2/3-IT1 neurons (Tukey-multiple comparisons: ***p-adjusted<0.001) and *Sod1^G86R^*mCSN (Tukey-multiple comparisons: ***p-adjusted < 0.001) compared to healthy controls and WT mice respectively, and a trend to an increase in patients’ L5-ET (Tukey-multiple comparisons : p>0.05) **i.** Violin plot showing module eigengene expression of the turquoise module in each of the seven major cell classes. The turquoise module is only significantly increased in excitatory neurons (ANOVA: F_group_= 26.67, p<0.0001, Tukey post-hoc: *p-adjusted = 0.035). **j.** Violin plot showing module eigengene expression of the lightyellow module in each of the seven major cell classes. The lightyellow module is significantly increased in both the excitatory neurons (ANOVA: F_group_= 26.67, p<0.0001, Tukey post-hoc: ****p-adjusted < 0.0001) and the inhibitory neurons (ANOVA: F_group_= 26.67, p<0.0001, Tukey post-hoc: ***p-adjusted < 0.001). **k.** Expression of the turquoise and lightyellow modules in *Sod1^G86R^* mCSN at presymptomatic (30 and 60 days) and symptomatic (90 and 105 days) stages (ANOVA: F_group_ = 45.25, p<0.001; Tukey post-hoc: WT vs *Sod1^G86R^* at 30d p=0.075; 60d p=0.27; 90d ***p<0.001; 105d p=0.089). **l.** Representative images of puromycine incorporation in mCSN (CTIP2-positive neurons in layer 5 motor cortex) from 90-day-old *WT* and *Sod1^G86R^*animals, upon intracerebroventricular injection. Scale bar=20µm. **m.** Dot plot showing the quantification of the puromycin incorporation in mCSN (n=5 *WT* and 6 *Sod1^G86R^*; two-tailed nested t-test; *p=0.0207).

Gene Ontology (GO) analysis of the turquoise module revealed a strong enrichment in RNA-binding genes such as *EIF4A2*, *VCP*, *HNRNPA0*, *SF3A3* and *TUBA1B* (**Fig. 2e**, **Supplementary Table 10**), and terms associated with mRNA translation, ribosome, and mitochondrial translation (FDR < 0.05). The average expression of the turquoise module eigengenes revealed a significant increase in human L2/3 IT-1 and mCSN, and a trend to an increase in human L5-ET (**Fig. 2f**, **Supplementary Table 11**). GO analysis on the lightyellow module highlighted terms related to ubiquitin activity such as *HECTD1*, *UHRF2*, *UBE2K*, and terms associated with translation initiation factor activity such as *EIF2B4*, *EIF3L*, *EIF2D*, *EIF2S1*, the latter encoding the EIF2-alpha protein, a master regulator of the endoplasmic reticulum stress response (ER stress) (**Fig. 2g**, **Supplementary Table 10**). Additionally, we observed a significantly increased expression of the lightyellow module eigengenes in human L2/3-IT-1 and mCSN and a trend in human L5-ET (**Fig. 2h**, **Supplementary Table 11**).

Since we identified a strong enrichment of mCSN DEG in a large population of excitatory neurons (**Fig. 1d**), we tested whether the increase of the turquoise and lightyellow module eigengenes would be conserved in human cell types other than L2/3-IT-1 and L5-ET. To do so, we aggregated all neuronal and non-neuronal human subtypes into seven major classes ^8^ and assessed the turquoise and lightyellow module eigengene expression across species and genotypes/conditions. Human excitatory neurons showed a significantly increased expression of the turquoise (**Fig.2i**, **Supplementary Table 12-13**) and lightyellow modules (**Fig. 2j**, **Supplementary Table 12-14**) while inhibitory neurons display a marked increase of the lightyellow module (**Fig. 2j**, **Supplementary Table 14**). Other cell types instead showed non-significant trends (**Fig. 2i,j**, **Supplementary Table 13-14**). The data suggest that the molecular mechanisms associated with vulnerable mCSN and L5-ET virtually affect all human cortical glutamatergic neurons.

To test the dynamics of the turquoise and lightyellow module eigengene regulation over time, we took advantage of our longitudinal mCSN neurons dataset. We observed an increased expression of both module eigengenes in *Sod1^G86R^* compared to WT mCSN (**Fig.2k**, **Supplementary Table 15**), that was significant at the symptomatic age of 90 days (adjusted-p = 0.089). To validate our findings, we tested whether mCSN displayed altered mRNA translation, as suggested by GO analyses of the two prioritized modules. *In vivo* puromycin assay ^9^ revealed a significant decrease of puromycin incorporation in motor cortex L5 CTIP2-potitive neurons from 90 day-old *Sod1^G86R^* mice compared to WT, indicative of a slower rate of protein synthesis in disease vulnerable neurons (**Fig. 2l-m**). Thus, our results identified a cross species transcriptomic footprint of disease vulnerable neurons associated to ER stress and related altered mRNA translation.

### CREB3 is a master regulator of excitatory neuron function and a novel protective factor from ALS

Since the turquoise module was strongly associated with ALS and preserved across species, we sought to identify potential upstream regulators. Intersection of the turquoise module genes with a list of TFs from the ENCODE project (**Fig.3a**, **Supplementary Table 16**) ^10^, identified 18 TFs (**Fig. 3b**). Hierarchical clustering based on TF-target gene expression across cell types, further distinguished a cluster of 6 TFs with a relatively high expression in L5-ET neurons (**Fig. 3b**), composed of 2 CREB family members, *CREB1*, *CREB3*, as well as *BRF2, DR1, ZHX2* and *SP2*. TFs-based WGCNA connectivity ranking further revealed the strong connectivity of CREB3 with the genes of the turquoise module (**Fig. 3b**, **Supplementary Table 17**).

**Figure 3:**
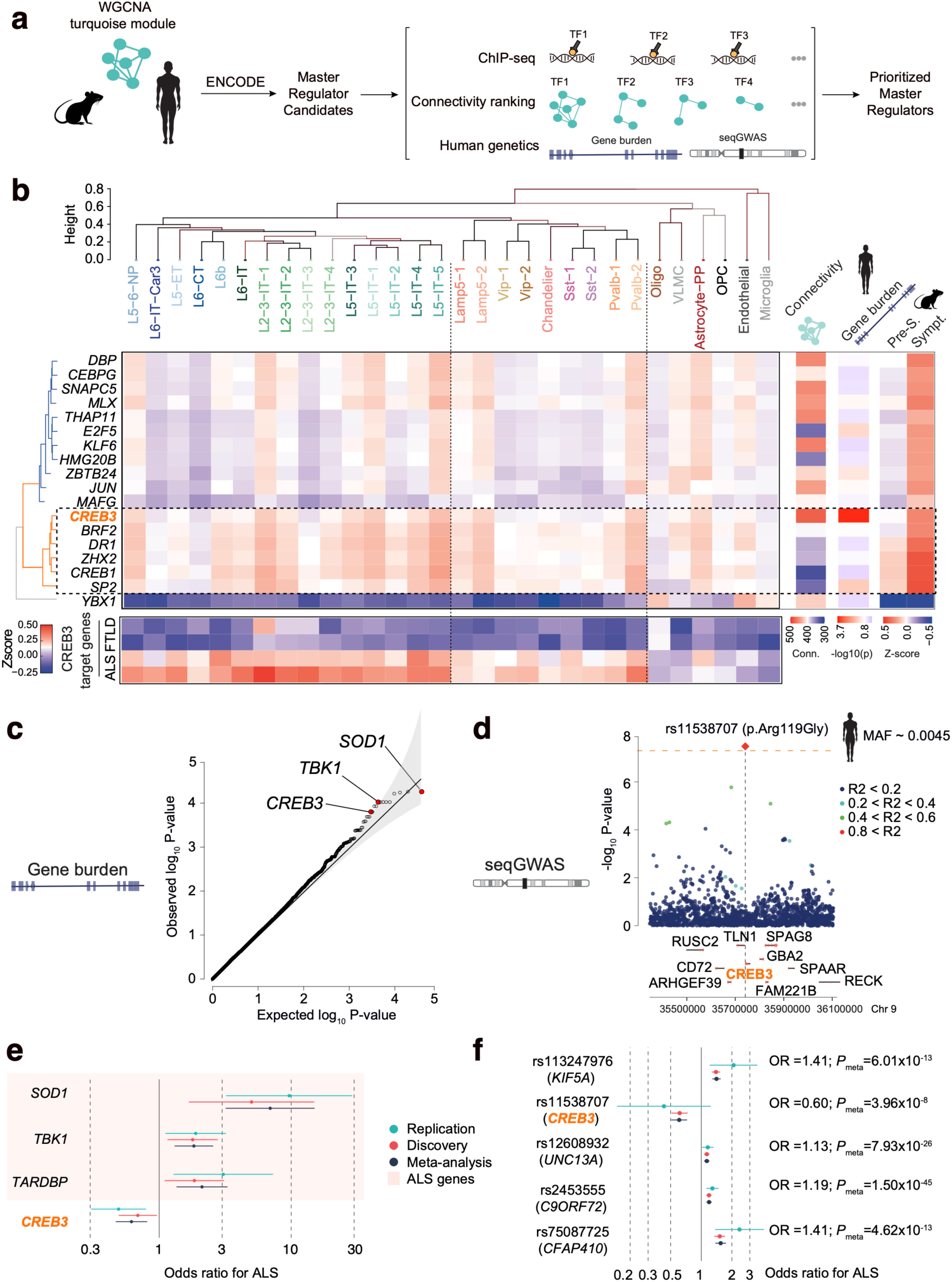
Integrative genetic analysis identifies the master regulator CREB3 as protective factor in ALS. **a.** Experimental design used to identify TFs upstream of the highly conserved and significant WGCNA turquoise module. Genes in the turquoise module were intersected with a list of transcription factors (TFs) from the ENCODE Consortium to identify potential master regulators. Each TF candidate was then further investigated using an integrative genetic and transcriptomic approach combining WGCNA-based connectivity, expression profile of the TF-target genes and burden of rare variants among identified TFs. **b.** Cluster dendrogram and heatmap showing the expression of the TF-target genes in each of the 30 DE human cell populations in ALS patients compared to healthy controls. Row clustering of TFs based on their target gene expression identifies two main clusters of TFs highlighted in blue and orange. The orange TFs cluster was further prioritized (dashed line rectangle) based on a higher expression level of their target genes in deep layer excitatory neurons (L5-ET). Right-sided heatmap showing WGCNA-based TF connectivity in prioritized cell populations (Fig. 1), association of missense rare variants in each TF with ALS risk, and expression profile of the target genes in mCSN at a presymptomatic (combined 30 and 60 days) and symptomatic (combined 90 and 105 days) stages. Bottom heatmap shows the expression profile of the prioritized TF *CREB3* in ALS and FTD patients compared to healthy controls. **c.** Quantile-quantile plot of the meta-analyzed gene burden of rare missense variants in a cohort of 1,018 ALS patients and 3,850 healthy controls showing *SOD1* gene as the top association, followed by other ALS known genes such as *TBK1*. **d.** Locus zoom plot showing the SNP (+/- 500KB) rs11538707 (R119G) association with ALS in the discovery cohort ^3^ and the replication of 1,018 ALS cases and 3,850 healthy controls. Orange dashed lines shows the genome-wide significant SNP at a p-value < 5^e-08^ and colored dots represent LD with the lead variant (red diamond). Forest plot of the gene burden association for known ALS genes such as *SOD1* (OR=5.23; 95%CI: 2.41-11.35; p=2.21^e-05^), *TBK1* (OR=1.79; 95%CI: 1.28-2.5; p=4.71^e-04^) and *TARDBP* (OR=1.97; 95%CI:1.26-3.07; p=2.1^e-03^) as well as the association of *CREB3* gene. Aggregation of rare missense variants in *CREB3* confers a reduced risk of ALS (OR=0.66 95%CI 0.51-0.87; p=2.9^e-03^). **f.** Forest plot showing four previously identified ALS associations on the *KIF5A*, *UNC13A*, *C9ORF72* and *CFAP410* genes, as well as the novel association of the missense variant rs11538707 (R119G) on *CREB3* with ALS risk (see Table 1).

To assess a putative role of CREB3 in ALS, we tested whether *CREB3* gene accumulates rare variants in ALS and/or healthy individuals, (**Fig. 3c**, **Supplementary Table 18**). We performed a rare missense variant burden analysis on a cohort of 1,018 ALS patients and 3,850 healthy controls which we meta-analyzed to the ALS Project Mine cohort (http://databrowser.projectmine.com/) ^11^. As expected, this revealed association in all known ALS genes such as *SOD1* (OR=6.93 [95%CI: 3.19-15.06]; P_meta_= 5.02e^-07^), *TBK1* (OR = 1.83 [95%CI:1.30-2.56]; P_meta_=2.21e^-04^) or *TARDBP* (OR = 2.11 [95%CI:1.35-3.29]; P_meta_=5.1e^-04^), conserved across cohorts, and highlighted *CREB3*, unravelling the protective effect of rare missense variants which replicated across independent cohorts (OR = 0.61 (95%CI:0.46-0.81); P_meta_=2.79^e-04^) (**Fig. 3c-e**, **Supplementary Table 18**). We then tested whether individual genetic variants could be associated with ALS risk through a sequenced-based genome-wide association study (seqGWAS) ^12^. A null logistic model was fitted on the cohort of 1,018 ALS cases and 3,850 controls. We observed a moderate inflation of the test statistics (λ_GC_ = 1.05), and linkage disequilibrium (LD) score regression yielded an intercept of 0.983 (s.e. = 0.0068), indicating that most of the inflation was due to the polygenic signal in ALS (LD score regression [LDSC]: ℎ2lhl2 = 0.69, s.e. = 0.0932). We were able to replicate most of the ALS GWAS variants identified by van Rheenen *et al.*, 2022 ^11^ (**Table 1**) such as *C9ORF72* (rs2484319, P_meta_= 1.50.10^-45^), *UNC13A* (rs12608932, P_meta_ = 7.93.10^-26^), *KIF5A* (rs113247976, P_meta_ = 6.01.10^-13^) and *CFAP410* (rs75087725, P _meta_ = 4.62.10^-13^) (**Fig. 3f**). In addition, we identified three novel loci through inverse-variance meta-analysis on chromosome 15 (rs12907456, P_meta_= 2.9.10^-08^), chromosome 17 (rs12907456, P_meta_= 2.75.10^-08^) and chromosome 9 (rs11538707, P_meta_= 3.96.10^-08^) (**Fig.3d-f, Table 1**). Most importantly, mapping rs11538707 to the human genome revealed a missense variant p.Arg119Gly (R119G) in *CREB3* which confers a 40% reduction in the risk of developing ALS (OR=0.61, 95%CI 0.51-0.73, **Fig.3f**). Overall, our cross-species transcriptomic analysis yielded a specific gene network associated with ALS from which *CREB3* appears as a major master regulator, and genetic data integration further identified *CREB3* as a protective factor in ALS.

### CREB3 regulatory network as a resilience marker in ALS

To determine if *CREB3* is actually causing altered gene expression in vulnerable neuronal populations, we performed a series of orthogonal approaches. First, we investigated the expression of *CREB3* in RNAseq data from mCSN. Both *CREB3* mRNA (**Fig.4a** FDR < *p < 0.05, **Supplementary Table 4**) and CREB3-target genes (**Fig. 4b**, **Supplementary Table 19**) were significantly upregulated in *Sod1^G86R^* mCSN. A strong up-regulation of CREB3-target genes was also observed in human excitatory and inhibitory neurons of the motor and frontal cortex from ALS patients, but not from FTLD patients (**Fig. 4c**, **Supplementary Table 20**), suggesting that CREB3 hyperactivity might be a marker of neuronal resilience in ALS. To confirm changes of *CREB3* expression at the cellular level, we performed fluorescent *in situ* hybridization and immunostaining on WT and *Sod1^G86R^*tissues at early symptomatic age or disease end-stage, respectively, which confirmed a significant increase of *CREB3* mRNA in L5 *Fezf2*+ neurons (**Fig. 4d**), and a significant increase of CREB3 protein in L5 CTIP2+ neurons of the motor cortex (**Fig.4e**). In a recent report, Moya and collaborators identified, among motor cortex L5 neurons of the *SOD1^G93A^*mouse model of ALS, disease vulnerable and disease-resistant subpopulations, respectively expressing *Gprin3* and *Colgalt2* ^13^. Leveraging these published TRAPseq data, we observed a significant increase of the turquoise module and of CREB3-target genes in disease vulnerable *Gprin3*-positive neurons compared to *Colgalt2*-positive neurons and whole motor cortex (**Supplementary Fig. 5a-c**). Finally, to test the extent of our findings in other ALS-vulnerable neurons, we investigated the recently published snRNAseq database of *SOD1^G93A^* spinal cord ^14^, and observed a significant upregulation of the turquoise module and of CREB3-target genes selectively in motoneurons, compared to astrocytes, microglia, endothelial cells, and other neuronal populations (**Supplementary Fig. 5d-f**). Overall, the data confirm that mCSN and human excitatory neurons display increased *CREB3* mRNA and protein expression, associated with the activation of CREB3 regulatory networks in ALS, and shows that this signature characterizes disease-vulnerable neuronal populations.

**Figure 4:**
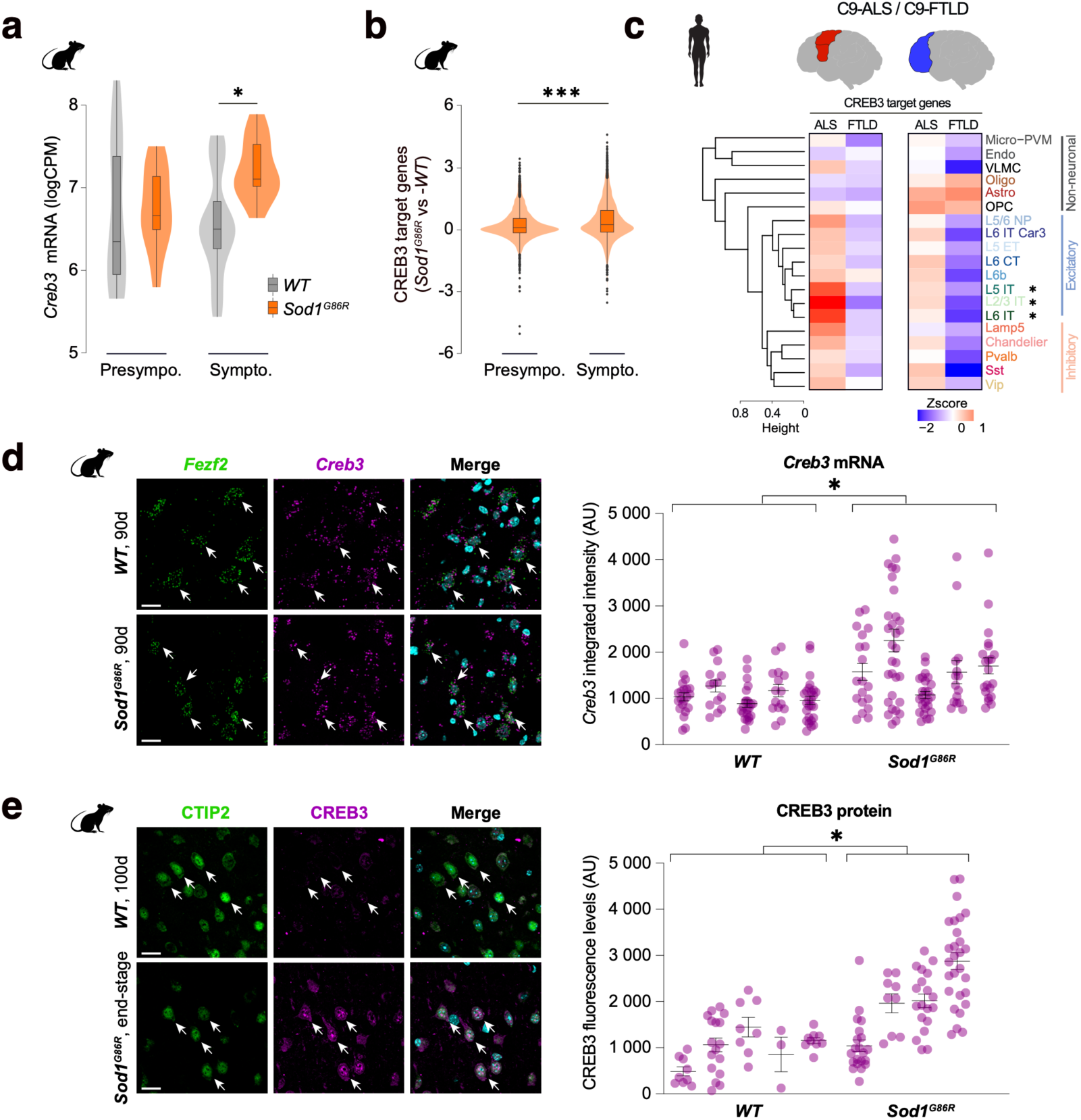
CREB3-target genes expression as a resilience marker in mouse and human neurons. **a.** Violin plot showing increased *CREB3* mRNA expression in mCSN of presymptomatic and symptomatic *Sod1^G86R^* mice (ANOVA: F_genotypes_ = 4.78, *p < 0.05, Tukey post-hoc: *p-adjusted = 0.029) **b.** Violin plot showing the expression of CREB3-target genes in motoneurons compared to other cell types in the spinal cord of *SOD1^G93A^* (Wilcoxon rank sum test: *p < 0.05, motoneurons vs all cell types). **c.** Heatmaps showing increased CREB3-target genes expression in the motor and frontal cortices of ALS patients versus FTDL patients. **d.** Representative images of the *Creb3* and *Fezf2* mRNA expression revealed by RNAscope in L5 of the motor cortex of 90-day-old WT and *Sod1^G86R^* mice (left, scale bar=20µm), and dot plot showing the quantification of *Creb3* probe integrated intensity in *Fezf2*+ neurons (n=1 female and 4 male *WT* mice, and n=2 female and 3 male *Sod1^G86R^* mice; two-tailed nested t-test; *p=0.0255). **e.** Representative images of CREB3 and CTIP2 immunoreactivity in L5 of the motor cortex of an end-stage *Sod1^G86R^* mouse and *WT* littermate (left, scale bar=20µm), and dot plot showing the quantification of CREB3 immunoreactivity in CTIP2+ neurons (n=1 female and 4 male *WT* mice, and n=1 female and 3 male *Sod1^G86R^* mice; averaged age at perfusion = 102 days for *WT*, 101 days for *Sod1^G86R^*; two-tailed nested t-test; *p=0.0207).

### CREB3 variant slows disease progression through gain of function

To further test whether increased transcriptional activity of CREB3-regulated genes would confer neuroprotection in ALS patients, we first identified, using CREB3-targeted chromatin immunoprecipitation and co-expression analysis, the CREB3 “regulon” which contains genes that would reflect CREB3 activity in tissues (**Fig. 5a**). We show that CREB3 regulon activity in the motor cortex (**Fig. 5b-c**) and in the blood of ALS patients (**Fig. 5d-e**) positively correlated with survival, and demonstrate an increased survival of ∼17 and ∼15 months respectively (**Fig. 5c, e**). The data highlight CREB3 regulatory network as a resilience marker in ALS.

**Figure 5:**
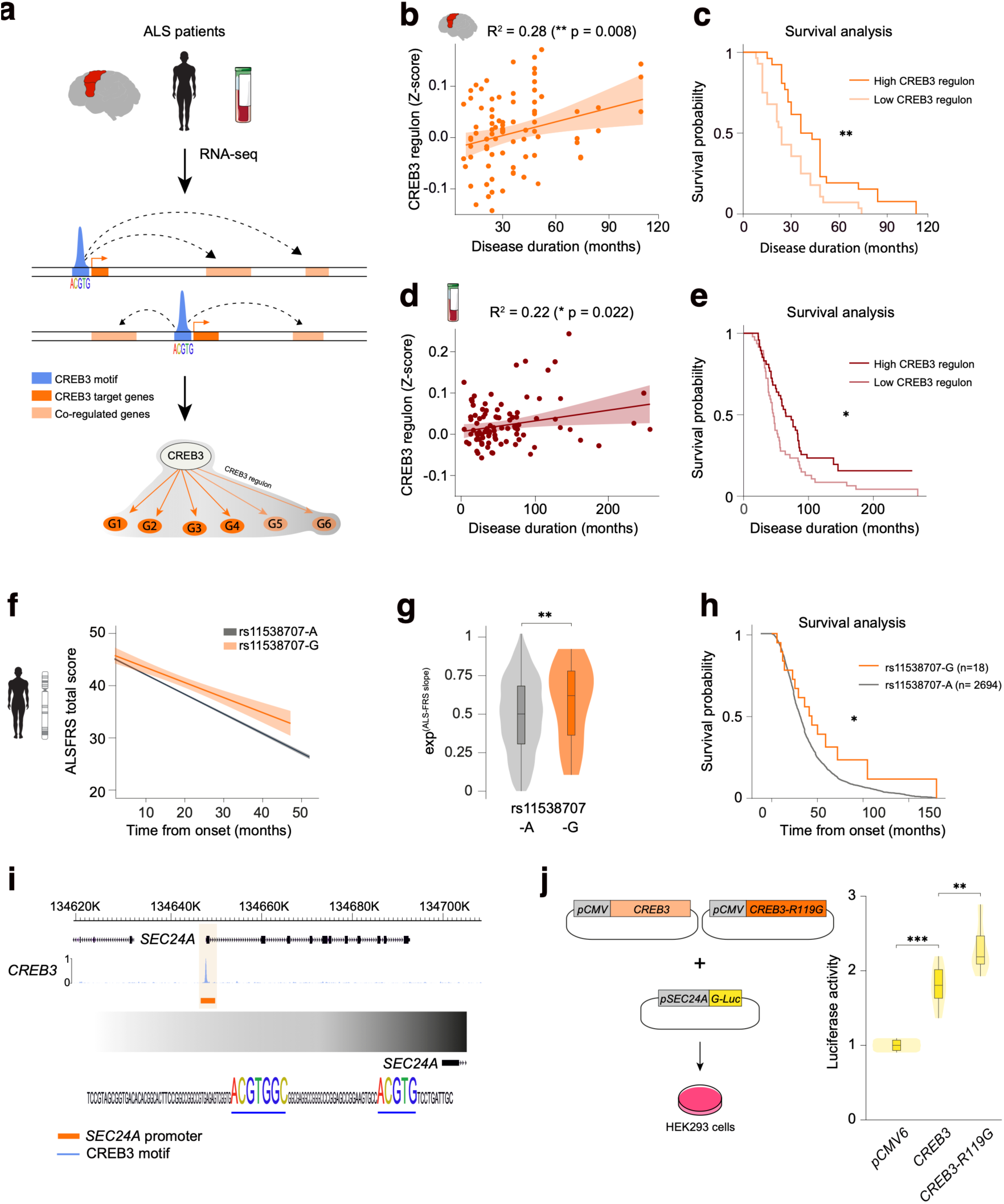
CREB3 gain of function is associated with a slower disease progression rate in ALS. **a.** Schematic identification of CREB3 regulon from RNAseq brain and blood data which encompasses CREB3-target genes identified through Chip-seq screening, and co-regulated genes identified through weighted gene co-expression analysis. **b.** Scatter plot showing significantly positive correlation between CREB3 regulon activity in the motor cortex of ALS patients and overall survival (Pearson R^2^ = 0.28; **p=0.008). **c.** Survival curve showing that ALS patients with higher brain CREB3 regulon activity have an overall longer disease duration (∼46 months) compared to ALS patients with lower CREB3 regulon activity (∼29 months) (Kaplan-Meier survival: Chi^2^ = 8.1; df = 1; **p = 0.004). **d.** Scatter plot showing significantly positive correlation between CREB3 regulon activity in the blood of ALS patients and overall survival (Pearson R^2^ = 0.22; *p=0.022). **e.** Survival curve showing that ALS patients with higher blood CREB3 regulon activity have an overall longer disease duration (∼77 months) compared to ALS patients with lower CREB3 regulon activity (∼62 months) (Kaplan-Meier survival: Chi^2^ = 5.5; df = 1; *p = 0.02). **f.** ALS functional rating scale (ALSFRS) progression in rs11538707 (R119G) carriers showing slower disease progression (mean = -0.69 pts / months) compared to non-carriers (-0.93 pts / months). **g.** ALS patient’s carriers of the rs11538707-G (R119G) variants show a significantly slower disease progression characterized by an attenuation of the functional decline compared to non-carriers (Wilcoxon rank test ALS-FRS slope: Bonferroni’s adjusted **p = 0.00183). **h.** ALS patients rs11538707 (R119G) carriers show a significantly extended disease duration (mean = ∼50.1 months compared to non-carriers (mean= ∼ 38.5 months) (Kaplan Meier: Chi-squared = 4.1, *p = 0.04). **i.** Genome browser Chip-seq track showing CREB3 binding on the canonical target gene promoter (light orange highlight) of *SEC24A* gene. **j.** Design of *CREB3* and *CREB3-R119G* expression vectors and *SEC24A-GLuc-ON* reporter plasmid transfected in HEK293 cells (left), and violin plots (right) showing increased *SEC24A* promoter activity in CREB3-transfected cells compared to control empty vector (Nested ANOVA: F_2-15_ = 45, p < 0.0001; Bonferroni’s test: ***p < 0.0001), further enhanced by the R119G missense mutation (Nested ANOVA: F_2-15_ = 45, p < 0.0001; Bonferroni’s test: **p = 0.0068).

Finally, we asked whether the R119G variant would also act as a disease modifier. Since R119G mutation acts as a protective factor, we observed a higher allele frequency in control (minor allele frequency = 0.47%) compared to patients (minor allele frequency = 0.21%), rendering difficult phenotyping characterization of rs11538707 carriers in ALS patients. However, leveraging a large cohort of whole-genome sequenced datasets, we show that carriers of the rare minor allele rs11538707-G have a slower progression rate characterized by a slower decline in ALS-FRS score over time (mean ALS-FRS slope = -0.69 pts/months) compared to rs11538707-A allele (mean ALS-FRS slope = -0.93 pts/months) (**Fig.5f-g**, **p<0.001). Importantly, the carriers of the rs11538707-G also show an extended lifespan with a mean disease duration of ∼ 50.1 months compared to the rs11538707-A carriers whose disease duration is 1 year shorter on average (mean disease duration ∼ 38.5 month) (**Fig.5h**, *p<0.05). Overall, this data confirms that the R119G mutation has a protective role in ALS and demonstrates that ALS patients that do carry the variant have a significant slower rate of progression.

Finally, to test whether the R119G mutation could modulate CREB3 transcriptional activity, we designed a reporter-based assay. We identified *SEC24A* as a canonical CREB3-target gene with multiple CRE-responsive elements in its promoter region (**Fig. 5i**), and generated a reporter plasmid by placing the luciferase gene downstream of the *SEC24A* promoter (**Fig. 5j**). Co-transfection of a wild type *CREB3* expression plasmid increased *SEC24A* promoter activity by a factor of 2, confirming *SEC2A4* as a *bona fide* CREB3-target gene (**Fig. 5j**). Co-transfection of a *CREB3* R119G expression plasmid further enhanced the *SEC24A* promoter activity by 48% (**Fig. 5j**). Thus, the *CREB3* R119G variant confers protection against ALS via a higher transcriptional activation of its target genes.

## Discussion

In this study, we tested the possibility that at least a subpart of genetic disease modifiers of ALS may directly influence the molecular pathways selectively activated in vulnerable neurons as the disease progresses, and concentrated on CSN for their selective vulnerability to ALS and their excitatory glutamatergic neuron identity ^3–5^. We drew inspiration from work carried out in the fields of evolution and development ^8,15–17^, and implemented comparative cross-species transcriptomics from post-mortem human snRNAseq and anatomically defined mCSN RNAseq from an ALS mouse model. We identified, as expected, a strong similarity between mCSN and human L5-ET neurons, which include the Betz cells ^18^, but also with human L2/3 IT-1, a population which, noteworthily, in non-human primates, is vulnerable to degeneration in late-stage Alzheimer’s disease ^19^. Using weighted gene co-expression analysis of mCSN, L2/3-IT1 and L5-ET neurons, we identified a conserved dysregulated gene network with a strong enrichment of genes associated with perturbed translational machinery and ER stress, a signature typical of neurodegeneration ^20^, and repeatedly reported in ALS models and patients ^21–25^. Whereas ER stress and UPR were expected in disease-vulnerable neuronal populations, we here demonstrate their presence across populations of excitatory neurons, indicating that, in ALS, there is no cell-type specific transcriptomic signature selective of disease-vulnerable neurons, but rather a conserved dysregulation of gene networks presents in broad populations of cortical neurons. By integrating single nuclei transcriptomics with human genetics and epigenetics, we further identified a set of 6 transcription factors whose target gene expression was increased in excitatory neurons of ALS patients. Importantly, using gene and variant based analysis, we further prioritized CREB3, whose target gene expression increased in excitatory neurons.

CREB3 belongs to a large bZIP family which regulates the expression of a large variety of genes involved in lipid metabolism, development, and protein secretion ^26^. Interestingly, a recent study in iPSCs-derived MN showed an increase of nuclear accumulation and phosphorylation of CREB1 (pCREB1^S133^) in ALS-patients-derived cultures in comparison with controls, followed by time-dependent depletion of pCREB1^S133^ which paralleled the synaptic impairment and preceded neuronal loss, indicative of an initial activation of a neuroprotective signaling that could not be sustained over time ^27^. Thus, one could speculate that, in stressed and disease-vulnerable long-range projection neurons, increased CREB3 activity may represent a protective mechanism allowing neurons to cope with disease-induced cellular changes, and contribute to slow-down the neurodegenerative process ^28^. In support of this argument, we show that ALS patients carrying the protective allele rs11538707-G have a significant longer disease duration compared to rs11538707-A carriers (∼51 vs ∼38 months), suggesting a neuroprotective effect of the p.Arg119Gly mutation through a CREB3 gain of function. Interestingly, previous studies have shown the critical role of regulated intramembrane proteolysis of ER-anchored transcription factor such as CREB3 in viral replication ^29^, and carriers of rs11538707-G protective allele were shown to have a significant increased risk of genital herpes infection ^30^ highlighting the role of CREB3 in antiviral response and immunity.

The most recent and largest GWAS in ALS to date identified numerous genes associated with ALS risk, but very few disease modifiers ^3^, such as the G4C2 expansion in the *C9ORF72* gene and *TBK1*, both associated with shorter survival ^3^. Yet, it is noteworthy that, similarly to *CREB3*, both genes have important functions in the immune response to pathogens ^31^ and are highly expressed in immune brain cells ^32^. More recently, using rare variant association analyses applied to whole genome sequencing data, Eitan *et al.* identified rare variants in the *IL18RAP* 3**′**UTR in non-ALS genomes, associated with a fivefold reduced risk of developing ALS ^33^. The authors further demonstrated that the variant *IL18RAP* 3′UTR attenuated NF-κB activity in microglia ^33^. It is therefore tempting to speculate that the immune-nervous system interaction could be the key driver of disease progression in ALS. Together with *IL18RAP* 3**′**UTR ^33^ and *KIFAP3* ^34^ that encodes a kinesin-associated motor protein, *CREB3* adds to the extremely short list of protective variants against ALS.

Overall, through integrative cell-type specific comparative analysis of mouse and human transcriptomics, combined to large scale genetic and epigenetic data, we identified CREB3 as a new protective factor from ALS, that confers neuroprotection through a gain of function mechanism. Future studies will be needed to understand the role of CREB3 in neurons. As of now, our data suggest that the differential cellular vulnerability of CSN is associated with their intrinsic properties of long-range projection neurons, and that boosting CREB3 activity could represent an interesting therapeutic strategy to prevent disease onset or slow-down disease progression.

## Materials and Methods

### mCSN RNAseq

#### Animals

All animal experiments were performed under the supervision of authorized investigators and approved by the local ethical committee of Strasbourg University (CREMEAS, agreements #00766, #1534 and #28521). BAC transgenic male mice with the G86R murine Sod1 missense mutation ^35^ were obtained from the animal facility of the Faculty of Medicine, University of Strasbourg. Non-transgenic age-matched male littermates served as controls. Mice received water and regular rodent chow ad libitum. *Sod1^G86R^* animals were followed daily, and disease progression was rated according to a clinical scale going from score 4 to 0, as we previously described ^36^.

#### Retrograde labelling of mCSN and FACS purification of mCSN and mCtl cells

25-day-old WT and *Sod1^G86R^* mice were deeply anesthetized with an i.p. injection of Ketamine (Imalgène 1000®, Merial; 120 mg/kg body weight) and Xylazine (Rompun 2%®, Bayer; 16 mg/kg body weight) and placed on a heating pad. A laminectomy was performed on C3-C4 cervical vertebrae and the animals were positioned below an injector (Nanoject II, Drummond Scientific, PA) mounted on a micromanipulator. A pulled glass capillary loaded with Green IX Retrobeads (Lumafluor) was used to puncture the dura and lowered to the dorsal funiculus. Five pressure microinjections of 23 nl were performed on each side of the dorsal funiculus. 30, 60, 90 105-day-old injected mice were deeply anesthetized with by an i.p. injection of Ketamine/Xylazine (120 mg/kg; 16 mg/kg) before decapitation. The brains were sectioned in a stainless-steel coronal brain matrix (Harvard Apparatus, MA), and 1 mm-thick sections were transferred under a fluorescence SMZ18 microscope (Nikon). Cortical layer 5 and layers 2-3 from same animals were microdissected from 4 coronal sections and collected in separate tubes filled with iced HABG (Hibernate A (BrainBits UK), B27, Glutamax (Gibco) and 0.1N NaOH). The microdissected tissues were enzymatically digested with 34 U/ml papain at 37°C for 30 min. Cells were mechanically dissociated by gentle trituration in iced HABG, filtered through 70 μm cell strainer (BD Falcon) and subjected to density centrifugation through a three-density step gradient of Percoll (Sigma, MO) as previously described ^37^. Upon centrifugation, the cell pellets were resuspended in cold 0.01 PBS and fixed with 70% EtOH for 30 min at 4°C. Fixed cells were centrifugated to eliminate EtOH and resuspended in 0.01 M PBS complemented with RNAse inhibitors (Promega). Microsphere-labelled mCSN were purified using the FACS Aria II (BD Biosciences), based on their fluorescence, size and granularity. Unlabelled control cells were purified by FACS, using the same size and granularity as for the mCSN. Approximately 2,000 CSN and exactly 2,000 control cells were collected from each adult mouse brain and used as individual biological replicates for RNA sequencing.

#### RNA sequencing

Full length cDNA was obtained using the SMART-Seq v4 Ultra Low Input RNA kit for Sequencing (Clontech, CA) according to the manufacturer’s instructions. 11 cycles of cDNA amplification were performed using the Seq-Amp polymerase. 600ng of pre-amplified cDNA were then used as input for Tn5 transposon tagmentation by the Nextera XT kit (Illumina) followed by 12 cycles of library amplification. Following purification using Agencourt AMPure XP beads (Beckman Coulter), the size and concentration of the libraries were assessed on an Agilent 2100 Bioanalyzer. The libraries were then loaded in the flow cell at a concentration of 3nM, clusters were generated by using the Cbot and sequenced on the Illumina HiSeq 4000 system as paired-end 2x50-base reads, following Illumina’s instructions. Raw reads were mapped to the mouse reference genome GRCm38 with STAR version 2.7.0 ^38^ and default parameters using Ensembl gene annotations (version 87). Gene-level abundance estimates were estimated using the option–quantMode geneCount in STAR. We filtered the lowly expressed genes wherein each gene was required to have at least 15 counts across all samples and used both exonic and intronic reads. The filtered set of genes was used for the PCA plot and differential expression analysis. Differential gene expression analysis was performed with the ARMOR workflow ^39^ and a cut off FDR value of 0.05 was set in both datasets.

#### Data repository

RNA-seq data have been deposited on the ArrayExpress database at EMBL-EBI under the accession number E-MTAB-7876: https://www.ebi.ac.uk/arrayexpress/experiments/E-MTAB-7876/

### Human single nucleus RNAseq (snRNAseq) data analysis and cluster annotation

#### SMART-seq v4

Raw read (fastq) files from the Allen Brain Atlas ^8^ were aligned to the GRCh38 human genome sequence (Genome Reference Consortium, 2011) with the RefSeq transcriptome version GRCh38.p2 (RefSeq, RRID SCR_003496, current as of 13 April 2015) and updated by removing duplicate Entrez gene entries from the gtf reference file for STAR processing as described previously ^8^.

#### 10× Chromium RNA sequencing (Cv3)

Raw FASTQ files were downloaded from Pineda et al., 2023 ^40^ and the Allen Brain Atlas ^8^ and aligned to the pre-mRNA annotated human reference genome GRCh38 using Cell Ranger v4.0 (10x Genomics, Pleasanton CA) except for substituting of the curated genome annotation used for SMART-seq v4 quantification. Introns were annotated as ‘mRNA’, and intronic reads were included to quantify expression. Quality control criteria were used as previously described ^8^, so that for Cv3, criteria were: more than 500 (non-neuronal nuclei) or more than 1,000 (neuronal nuclei) genes were detected and doublet score was less than 0.3.

#### Clustering of snRNA-seq data

Nuclei were grouped into transcriptomic cell types using an iterative clustering procedure ^41^. Read counts were summed, and log_2_-transformed expression was centered and scaled across nuclei. Clusters were identified with Louvain community and pairs of clusters were merged if either cluster lacked marker genes. Clustering was applied iteratively to each subcluster until clusters could not be further split and robustness was assessed by repeating iterative clustering 100 times for random subsets of 80% of nuclei. Consensus clusters were defined by iteratively splitting the co-clustering matrix. The clustering pipeline is implemented in the R package scrattch.hicat v0.0.22 with marker genes defined using the limma package; the clustering method is provided by the ‘run_consensus_clust’ function (https://github.com/AllenInstitute/scrattch.hicat). Clusters were curated based on quality-control criteria or the expression of markers genes ^8^.

To establish a set of human consensus cell types across all three datasets, we performed a separate integration of snRNA-seq technologies on the major cell classes (glutamatergic, GABAergic, and non-neuronal) as described previously ^8^.

Each expression matrix was log_2_(CPM + 1) transformed then placed into a Seurat object and variable genes were determined by down sampling each expression matrix to a maximum of 300 nuclei per scrattch.hicat-defined with n set to 20, to generate a list of up to 20 marker genes per cluster. The union of the Cv3, Cv3-ALS and SSv4 gene lists were then used as input for anchor finding, dimensionality reduction, and Louvain clustering of the full expression matrices. Louvain clustering was performed to over cluster the dataset to identify more integrated clusters than the number of scrattch.hicat-defined clusters. For instance, glutamatergic neurons had 30, 47 and 48 scrattch.hicat-defined clusters, 110 overclustered integrated clusters, and 38 final human consensus clusters after merging for Cv3 and SSv4 datasets, respectively. To merge the over clustered integrated clusters, up to 20 marker genes were found for each cluster to establish the neighborhoods of the integrated dataset. Clusters were then merged with their nearest neighbor if there were not a minimum of ten Cv3 and two SSv4 nuclei in a cluster, and a minimum of 4 DEGs that distinguished the query cluster from the nearest neighbor.

#### Cell-type specific differential expression

Differentially expressed genes (DEGs) for a given species were identified by using Seurat’s FindAllMarkers function with a Wilcox test and comparing each cluster with every other cluster under the same subclass, with logfc.threshold set to 0.7 and min.pct set to 0.5. The union of up to 100 genes per cluster with the highest avg_logFC was used. The average log_2_ expression of the DEGs was then used as input for the build_dend function from scrattch.hicat to create the dendrograms.

Cell type-specific pseudo-bulk differential gene expression (DGE) groups were built based on hierarchical clustering and Euclidean distance between each cluster leading to 30 DGE cell groups (see Fig. 2). For each of the DGE cell group, differential expression analysis was performed as described in the ARMOR workflow ^39^ for sufficiently abundant cell types using age, sex, and disease group as design covariates and gene-wise single-cell-level variance as weights for the linear model.

### CSN gene signature in human snRNAseq

A geneset of GFP-positive cells was built based on DGE analysis of CSN vs and control cells in mouse. Our final set of ∼50 genes was used as an input to AUcell ^42^ to evaluate the distribution of AUC scores across all the cells and explore the relative expression of the signature. The function AUCell_exploreThresholds() was used to determine the minimum AUC values where cells are considered to express the CSN geneset. For each cluster, the proportion of cells expressing the CSN geneset was calculated.

To identify overlap between mouse and human snRNAseq gene signatures, cell-type specific DGE genes were intersected with mCSN DEG genes, and significant overlap was calculated using an hypergeometric test with the phyper() function in R.

### Cross-species consensus weighted-gene coexpression network analysis (WGCNA)

To facilitate comparison across species, mouse gene identifiers were re-annotated with human Ensembl gene orthologs using biomaRt, an R interface with the Biomart database (www.biomart.org) ^43^. Only identifiers that were common to both human and mouse meta-sets were retained. A consensus network represents a single network arising from multiple sources of data constructed from the weighted average of correlation matrices from both the human and mouse in this study. By definition, consensus modules are the branches of a clustering tree developed from a consensus gene dissimilarity, comparable to the single-network approach; consensus modules contain genes that are closely related in both networks, i.e., the modules are present in both networks. After scaling the network (consensus scaling quantile = 0.2), a threshold power of 14 was chosen (as the smallest threshold resulting in a scale-free R^2^ fit of 0.9) and the consensus topological matrix was created as follows : consTOM <- consensusTOM(multiExpr,checkMissingData = TRUE,maxBlockSize = Inf, randomSeed = 12345, corType = “pearson”, maxPOutliers = 0.05, quickCor = 0, pearsonFallback = “individual”, power = 14, networkType = “signed”, TOMType = “signed”, networkCalibration = “full quantile”, calibrationQuantile = 0.95,sampleForCalibration = TRUE, sampleForCalibrationFactor = 5000). The consensus tree was then built and the modules identified with the function cutTreeDynamic() with the default parameters and the minModuleSize (=30). This approach identified 36 modules for which correlation to phenotype (ALS) and adjusted-pvalue (FDR) were calculated and intersected between mouse and human to identify conserved directionality and association. This approach led to the identification of a final set of 20 modules. Module eigengenes (MEs) were used for module–trait association analysis, differential eigengene network analysis, and for differential gene expression analysis. Difference in expression across trait groups was tested using a Kruskall–Wallis one-way analysis of variance. A gene’s module membership (k _ME_) is defined as the Pearson correlation between each gene and each ME; genes with high k _ME_ values were considered “hub” genes and were highly co-expressed within a subnetwork. Module preservation statistical tests were used to assess how well network properties of a module in one reference data set were preserved in a comparator data set (modulePreservation function in WGCNA). Preservation statistics are influenced by a number of variables (module size, network size, etc). A composite preservation Z-score (Z _summary_) was used to define preservation relative to a module of randomly assigned genes where values 5 < Z < 10 represent moderate preservation, while Z > 10 indicated high preservation. Genes in each network module were characterized using EnrichR (version 1.2.5) ^44^, and we considered a term to be significant for FDR < 0.05. Genes network modules were constructed using metacells transformed data aggregated by cell class. For glutamatergic metacells matrices, we calculated signed kMEs for each gene and each module identified in the WGCNA analysis. The topological matrix was filtered to contain selected hub genes (2 gene/module) and was used an input to UMAP dimensionality reduction. Finally, each dot size (genes) was scaled to the signed kMEs for the corresponding module.

#### Violin plot of eigengenes expression

For each module and corresponding cell type, module eigengenes was calculated in control and ALS conditions using the moduleEigengenes() function. Comparison across conditions was calculated using a Kruskall–Wallis one-way analysis of variance and considered significant when FDR < 0.05.

#### ENCODE Chip-seq transcription factors

A list of 281 transcription factors and their bed files were downloaded from the ENCODE consortium (https://www.encodeproject.org/). Bed files were intersected with TSS annotations from the GRCh38 version and peaks were annotated to the closest TSS genes.

A final list of 278 TFs was intersected with genes identified in the WGCNA turquoise module, which lead to a final set of 18 TFs candidates. For each of TFs candidate, target gene Z-scores were average for each cell type and cluster using the hclust() function in R. CREB3 target gene Z-scores were averaged across cell types and compared between FTD and ALS patients.

### Genetic data processing and association

#### Association testing and meta-analysis

Whole genome sequencing from 1,345 ALS patients and 3,860 controls were processed as described above. Duplicate individuals were removed (king-cutoff = 0.084). Population structure was assessed by projecting 1,000G principal components (PCs) and outliers from the European ancestry’ population were removed (> 4 SD on PC1-4). Finally, samples in common between the individual genotype data and van Rheenen’s study ^3^ were identified using the checksum program id_geno_checksum and were removed from our analyses. In total, 1,018 ALS cases and 3,850 controls pass quality check analysis and were used for rare variant burden and genome-wide association analyses (GWAS). After quality control, a null logistic mixed model was fitted using SAIGE with principal component (PC)1–PC10 as covariates. The model was fit on a set of high-quality (INFO > 0.9) SNPs pruned with PLINK 2.0 (‘–indep-pairwise 50 25 0.2’) in a leave-one-chromosome-out scheme. Subsequently, a SNP-wise logistic mixed model including the saddle point approximation test was performed using genotype dosages with SAIGE. To assess any residual confounding due to population stratification and artificial structure in the data, we calculated the LDSC intercept using SNP LD scores calculated in the HapMap3 CEU population.

#### Genic burden association analyses

To aggregate rare variants in a genic burden test framework we used the method described in the ALS 2021 GWAS ^3^. In short, a variety of variant filters was applied to allow for different genetic architectures of ALS associated variants per gene as was used previously ^45,46^. In summary, variants were annotated according to allele-frequency threshold (MAF < 0.01 or MAF < 0.005) and predicted variant impact (“missense”, “damaging”, “disruptive”). “Disruptive” variants were those variants classified as frameshift, splice-site, exon loss, stop gained, start loss and transcription ablation. “Damaging” variants were missense variants predicted to be damaging by seven prediction algorithms (SIFT, Polyphen-2, LRT, MutationTaster2, Mutations Assessor, and PROVEAN). “Missense” variants are those missense variants that did not meet the “damaging” criteria. All combinations of allele frequency threshold and variant annotations were used to test the genic burden on a transcript level in a Firth logistic regression framework where burden was defined as the number of variants per individual. Sex and the first 20 principal components were included as covariates. All ENSEMBL protein coding transcripts for which at least five individuals had a non-zero burden were included in the analysis. Meta-analysis was performed using an inverse variance weighted method ^47^.

### Validations on mouse tissues

#### Puromycin administration

Six 90-day-old female *Sod1^G86R^* mice and five WT female littermates were anaesthetized with 2% isoflurane / 98% air, and two intracerebroventricular injections (one per hemisphere) were performed in the lateral ventricles to deliver 2.07µl of puromycin (Sigma, P7255) diluted in saline at 25µg/µl as previously described ^9^. Animals were allowed to recover from surgery and sacrificed and perfused one hour later. Their brains were collected and processed for immunofluorescence.

#### Immunofluorescence

Mice were euthanized with an overdose of pentobarbital sodium and phenytoin sodium (120 mg/kg) and transcardially perfused with cold 0.01 M PBS, followed by cold 4% PFA in 0.01 M PBS. Brains were dissected and post-fixed overnight in 4% PFA. Brains were cut coronally in 40μm-thick sections on a vibratome (Leica Biosystems). To reveal CREB3 expression, brain sections were first heated at 80°C in 10mM citrate buffer for 30 min. This step was not used to reveal puromycin. Brain sections were incubated 1h in blocking solution (8% goat serum, 0.3% BSA, 0.3% Triton in PBS), and 48 to 72h at 4°C with primary antibodies. Sections were rinsed, incubated 2h with the secondary antibodies, rinsed and mounted in Prolong Diamond mounting medium (Invitrogen, # P36970). The primary antibodies used were: mouse@puromycine (DSHB, #PMY-2H4; 1/100), rat@CTIP2 (Abcam, #Ab18465; 1/100) and rabbit@CREB3 (Aviva Systems Biology #OAAN03577, 1/50). Secondary antibodies were from the Alexa series (Life Technologies; 1/1000).

#### RNAScope

Upon fixation as described above, brains were cryoptotected and cut coronally in 14μm-thick sections on a cryostat (Leica). RNAscope® Multiplex Fluorescent Reagent Kit v2 (Advanced Cell Diagnostics, #323100) was employed according to the manufacturer’s instructions, using probes targeting Creb3 and Fezf2 (Advanced Cell Diagnostics #1263611-C3 and #313301-C2). Briefly, sections were incubated with 1X PBS for 5min to wash out OCT, baked for 30min at 60°C, post-fixed with 4% fresh PFA for 90min, sequentially dehydrated with increasing concentrations of ethanol, baked again for 30min at 60 °C, and incubated for 10min in RNAscope™ Hydrogen Peroxide Reagent at RT. Target retrieval was performed for 5min in a steamer, and RNAscope™ Protease III Reagent was applied for 15min at 40°. Slides were then hybridized with target probes for 2h at 40°C in a hybridization oven (Boekel). Slides were stored with 5X Saline Sodium Citrate overnight at RT and signals were amplified using amplifiers and horse radish peroxidases from the reagent kit and TSA fluorophores (TOCRIS, # 7526 and # 7527). Samples were mounted in Superfrost® Plus slides (VWR, #631-0108) using ProLong™ Diamond Antifade Mountant medium (Invitrogen, #P36965).

#### Image acquisition and quantification

Images were captured at 63X using an AxioImager.M2 microscope equipped with a structured illumination system (Zeiss) and a high-resolution B/W camera (Hamamatsu), and run by the ZEN 2 software (Zeiss). Image analyses were performed with ImageJ (NIH). Modal Grey Value of puromycin and CREB3 signals was measured in CTIP2+ neurons in layer 5 of the mouse motor cortex, and Creb3 integrated density was measures in Fezf2+ cells.

### Dual luciferase assays

HEK-293 cells were cultured in Dulbecco’s modified Eagle’s Medium (DMEM), 10% foetal bovine serum and 1% penicillin-streptomycin at 37°C in 5% CO2. Cells were plated in a 6-well plate and transfected 24 h after plating in DMEM + 0.1% foetal bovine serum and 1% penicillin-streptomycin using Lipofectamine 2000 (Invitrogen). 800 ng of dual luciferase reporter containing CREB3 response element was co-transfected with 2400 ng of pCMV6, pCMV6-CREB3 or pCMV6-CREB3-R119G plasmid. Dual luciferase assays were performed 24h after transfection as described by the manufacturer (Promega Secrete-Pair Dual Luminescence Assay Kit # LF033).

### Statistics

Immunofluorescence and RNAScope data are represented as nested scatter dot plots, and were analyzed using two-tailed nested t tests performed on Prism 6 (GraphPad). Results were considered significant when p<0.05. For the luciferase assay, transfected cells were run as duplicate for a total of 6 independent experiment. A linear mixed-model was fitted to integrate the technical replicate as random effect followed by a Bonferroni’s test for multiple comparison. Results were considered significant when adjusted-p < 0.05.

## Consent for publication

All authors read and approved the publication of this manuscript.

## Authors’ contributions

CR and SM conceptualized the study. SM performed transcriptomic and genetic analysis. CM and MF generated mouse RNAseq data. SM, CK and AMP analyzed mouse RNAseq data, and SM analyzed human snRNAseq data and genetic data. MHG, GST, SDG, CG and AB performed immunofluorescence and RNAscope validations, with image analyses tools designed by PK. CS performed the luciferase assays designed by LD. LD and SM established the needed collaborations with geneticists. BT, LF and AC provided the whole genome sequencing data of ALS patients and controls from The Netherlands, Germany and Italy respectively, as well as clinical data. The manuscript was drafted by C.R and S.M and reviewed and accepted by all authors.

## Acknowledgements

The authors are thankful to the ENCODE and Project Mine Consortia which generated part of the datasets used in this study. They thank Claudia De Tapia, Annie Picchinenna and Marie-José Ruivo for technical support.

## Data availability

Mouse CSN RNAseq data have been deposited in the ArrayExpress database at EMBL-EBI under accession number E-MTAB-7876. Human single-nuclei RNA-seq are available through the Gene Expression Omnibus repository under the accession code: GSE174332 and GSE219281. Post-mortem brain RNAseq from ALS patients are available through the target ALS consortium (https://www.targetals.org/research/funded-consortia/) with restricted access to authorized researcher. Blood RNA-seq dataset is publicly available and can be accessed at NCBI Gene Expression Omnibus (GEO) under the accession code: GSE234297. Human ALS whole genome sequencing is available to authorized researcher through dbGaP under the accession: phs001963.v2.p1.

